# Automatic computational classification of bone marrow cells for B cell pediatric leukemia using UMAP

**DOI:** 10.1101/2025.03.03.641130

**Authors:** Ana Niño-López, Álvaro Martínez-Rubio, Rocío Picón-González, Ana Castillo Robleda, Manuel Ramírez Orellana, Salvador Chulián, María Rosa

**Affiliations:** Department of Mathematics, University of Cádiz, Puerto Real, Spain; Biomedical Research and Innovation Institute of Cádiz (INiBICA), Puerta del Mar University Hospital, Cádiz, Spain; Oncohematology Unit, Niño Jesús University Children’s Hospital, Madrid, Spain; Foundation for Biomedical Research Niño Jesús University Children’s Hospital, Madrid, Spain; Health Research Institute La Princesa, Madrid, Spain

**Keywords:** Acute Leukemia, Automation, UMAP, Flow Cytometry

## Abstract

B Acute Lymphoblastic Leukemia (B-ALL) accounts for approximately 80% of pediatric leukemia cases. Despite treatment advances, 15–20% of children experience relapse, highlighting the need of improved monitoring of patients and novel strategies leading to successful therapies.

Flow Cytometry is an essential technique for measuring residual disease and guiding treatment. However, traditional manual gating limits its efficiency. In recent years, computational tools have been integrated to enhance these clinical processes but many mathematical techniques are underexploited. Particularly, Uniform Manifold Approximation and Projection (UMAP), together with Machine Learning, provide promising approaches for analyzing large datasets. Mathematical tools and artificial intelligence offer new perspectives on these health problems, beyond the usual approach in biomedicine.

We have exploited 234 samples from 75 B-ALL patients to develop an artificial intelligence-based algorithm that can improve patient classification and therapy decisions in different patient cohorts. This implies an advancement on the routine manual analysis of the disease progression, as we identify key subpopulations automatically, distinguishing patients’ bone marrow regeneration patterns, thus improving the prediction and prognosis of the disease.

**MSC Classification:** 68-04, 68T09, 90C90, 92B05

**Abbreviations:** A list of abbreviations used throughout this work can be found in Appendix A.

## 1 Introduction

Acute lymphoblastic leukemia (ALL) is the most common childhood cancer (25%) and B-ALL accounts for 80% of all ALL in pediatric age. Despite improvements in treatments in recent decades, around 15-20% of B-ALL still experience a relapse (Velasco et al., 2023). One of the challenges remains identifying patients at high risk of relapse. Currently, patients diagnosed with ALL receive treatment according to protocols that adapt chemotherapy intensity to the estimated risk of relapse. Classification into these risk groups is based on biological characteristics of the patient and the disease, such as age, number of blasts at diagnosis, cytogenetics, molecular biology, and the response to treatment. This response is assessed by quantifying the levels of minimal residual disease (MRD), that is, the amount of disease present in a bone marrow aspirate obtained at specific times during treatment. Thus, for example, the Spanish protocol for pediatric ALL (SEHOP-PETHEMA-2013) quantifies the levels of MRD at days +15, +33 and +78, to make a clinical decision.

In particular, Flow cytometry (FC) is the technique of choice to perform cells quantification. This methodology has become a very powerful tool in clinical fields (Ng et al., 2024, Li, 2022). This technique to analyze patients samples can lead us to different studies to improve its use and improve the current utilities to expand the information obtained thanks to cytometry (Robinson et al., 2023). There are, in fact, works which combine genetic studies with computational tools (Tomar et al., 2019, Duan et al., 2022, Cheng et al., 2024, Jiwani et al., 2023) to analyze leukemic samples. Other works show how FC techniques allows doctors to detect the leukemic clone and distinguish a healthy bone marrow from a pathological bone marrow (Björklund et al., 2003, Sharma et al., 2023, Lebecque et al., 2024), using different computational techniques to manage the available information, yet manual gating is involved in the last steps from the whole process. However, and even if these techniques have been improved through the years (Velasco et al., 2023), to our knowledge there is not an automatic selection of leukemic cells or any automatic computational comparison between healthy and leukemic bone marrows.

Mathematical and computational tools can be combined and used along with data available to obtain new results, and it is increasingly common to find works related to the combination of these techniques (Ujas et al., 2023, Saeys et al., 2016, Stolarek et al., 2022, Seifert et al., 2023). Dimensionality reduction is a problem that has been studied for years (Carreira-Perpinán, 1997, Van Der Maaten et al., 2009). In particular, many studies analyze the application of these techniques related to cluster cell data analyzing the information with dimensional reduction, UMAP is highlighted in several works which compare different techniques (Becht et al., 2019, Allaoui et al., 2020). UMAP is faster and more meaningful in relation to organization of clusters obtained. As indicated, there are works related to find the disease in the sample applying this dimensionality reduction method (Weijler et al., 2022) but current works related to UMAP and FC focus on identifying the leukemic clone and always using manual techniques. We study the bone marrow samples, applying UMAP as a dimensionality reduction technique with a new perspective for analyzing the samples. We work on filling the gap in this area, by studying the bone marrow beyond this manual identification and thus automatically classifying cell groups. In this sense, we propose the creation of a general pipeline in hematological pathologies to automatically compare with bone marrow standard populations.

This proposal is motivated by recent advances in the field, where machine learning and dimensionality reduction techniques have shown promising results in clinical flow cytometry. For instance, Reiter et al. (2019) proposed an automated GMM-based approach to quantify measurable residual disease (MRD) in pediatric B-ALL, validated across 337 samples with high concordance to expert analysis and superior performance compared to other automated methods. Chulián et al. (2020) applied computational models to assess the prognostic relevance of immunophenotypic markers, showing that low CD38 expression at diagnosis correlated with increased relapse risk. More recently, Shopsowitz et al. (2024) introduced MAGIC-DR, which integrates UMAP and XGBoost to detect MRD in AML without manual gating, achieving excellent agreement with standard methods (AUC ≃ 0.97) and identifying undetected abnormal populations. Similarly, Driessen et al. (2024) used a non-supervised autoencoder with a classifier to detect malignant blasts at the single-cell level in pediatric AML, reaching over 90% accuracy and revealing developmental and immunophenotypic shifts between diagnosis and relapse—particularly in KMT2A-rearranged cases. Park et al. (2025) compared UMAP and t-SNE for MRD detection in leukemias and lymphomas, reporting over 95% concordance with manual analyses and sensitivity down 10^−5^. Lastly, Pedreira et al. (2025) benchmarked five classification algorithms for mature B-cell neoplasms, showing that CA-vSD and SVM achieved the best accuracy (up to 90.6%), although with trade-offs in diagnostic coverage.

With the aim of understanding the bone marrow composition, we focus on reducing the amount of information and simplifying current techniques to identify cells. In this work, we have samples analyzed by FC from patients diagnosed with B-ALL at different time points. Therefore, based on the tree structure of the hematopoiesis, in addition to identifying bone marrow cell subpopulations, we focus on finding markers that mainly distinguish B lymphocytes (Bendall et al., 2014, Fasano and Sequeira, 2017) according to their maturation states. To address this, we propose a semi-automated pipeline that (1) identifies both regenerated and leukemia-invaded bone marrow samples, and (2) monitors the dynamics of B-cell lineage recovery, enabling the extraction of immunophenotypic patterns that stratify patient cohorts and may inform relapse risk. Thus, the final aim of this study is to manage B-ALL data obtained by FC and to optimize current analysis techniques to improve precision and time through process automation.

## 2 Material and methods

### 2.1 Patients

In this study, 116 patients diagnosed with B Acute Lymphoblastic Leukemia (B-ALL) from Hospital Niño Jesús, Madrid (HNJ), have been considered. These patients, all in pediatric ages (1 − 19 years old), were diagnosed, treated and monitored according to SEHOP-PETHEMA 2013 protocol. The diagnoses related to these patients are dated between September 2013 and January 2022. Table 1 summarizes the clinical data of these patients, along with the features evaluated through integrated diagnosis and considered distinctive for each risk group according to the SEHOP-PETHEMA protocol.

**Table 1.** Characteristics of HNJ patients. Detailed information about patients included in this work.

| Feature | Values |  |
| --- | --- | --- |
| Sex | no. | % |
| Male | 56 | 48.3 |
| Female | 60 | 51.7 |
| Age at diagnosis | years |  |
| Median | 4 |  |
| Range | 1 – 16 |  |
| Long-term outcome | no. |  |
| Relapse | 25 | 21.6 |
| Non-relapse | 91 | 78.4 |
| Immunophenotype | no. | % |
| Common | 108 | 93.1 |
| Pre-B | 5 | 4.3 |
| Pro-B | 3 | 2.6 |
| Mixed | 0 | 0.0 |
| Bone marrow blasts at diagnosis | % |  |
| Median | 84.0 |  |
| Range | 3.0 – 99.0 |  |
| Karyotype | no. | % |
| Hyperdiploidy (> 50) | 18 | 15.5 |
| Normal (40–50) | 69 | 59.5 |
| Hypodiploidy (< 40) | 0 | 0.0 |
| No metaphases | 27 | 23.3 |
| Chromosomal alterations | no. | % |
| ETV6/RUNX1 | 29 | 25.0 |
| TCF3/PBX1 | 5 | 4.3 |
| MLL rearrangement | 2 | 1.7 |
| BCR/ABL1 | 3 | 2.6 |
| .fcs files | no. |  |
| At diagnosis | 116 |  |
| +33 days | 83 |  |
| +78 days | 82 |  |
| > +78 days | 9 |  |

#### Ethics statement

This retrospective study was designed in accordance with the Declaration of Helsinki, under the LLAMAT Project protocol (2018). Approval was granted by the institutional review boards (IRB) of the Hospital Niño Jesús, Madrid. Patients and/or their legal guardians signed a written, formal informed consent to participate in the study. Personal information was anonymized.

### 2.2 Flow cytometry data

The collected bone marrow samples are analyzed using flow cytometry, obtaining *.fcs* files to manage. This technique allows each cell in the sample to be studied separately, associating a value related to the intensity for each immunophenotypic biological marker (IPT). This is how samples are analyzed to identify the type of patient in the laboratory to assign each patient the corresponding treatment according to the protocol. Thus, the different types of ALL just like the subgroups of B-ALL (Ortunõ Giner and Orfao, 2002), are defined depending on the information obtained from flow cytometry.

Cytometers work through the reception of the fluorescence emitted by the cells regarding the chemical products used. FC consists of analyzing each single cell to associate an intensity value to each cellular immunophenotypic characteristic (IPT), which characterizes the cell type (Figure 1) and functions, as well as, for the case of leukemia patients, several risk groups (Orfao et al., 2019, Van Lochem et al., 2004). In this work, the main IPT markers to distinguish groups of cells in bone marrow are CD19, CD66 and CD45. It is important to highlight that there are four maturation states of B lymphocytes which are Pro-B, Pre-B, Transition and Mature, where intensities of IPT markers CD10 and CD45 allow us to classify them. All information from FC technique is collected as a data matrix in which each row corresponds to each of the cells and each column to each IPT marker. Due to the dimensions in which the data is presented, laboratory clinicians in the hospital have computer tools to be able to analyze the samples in two dimensions, comparing two markers on the x and y axes (Van Lochem et al., 2004, Wood, 2015). Data collection through FC technique involves a large amount of data that must be managed.

**Fig. 1.**
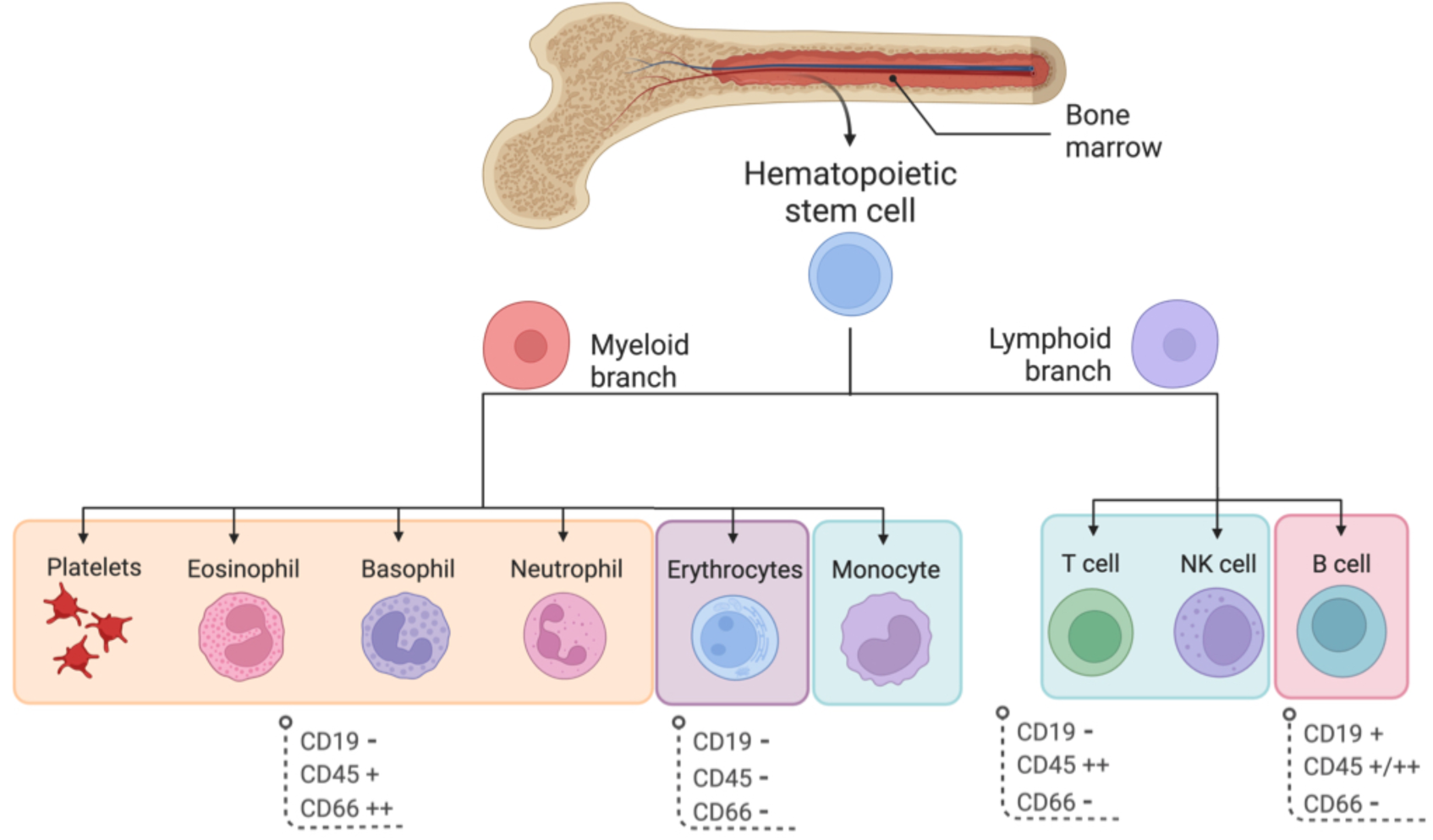
Biomarkers associated to different types of cells in bone marrow. We focus on CD19, CD45 and CD66 as IPT markers to distinguish four main subgroups of cells (Orfao et al., 2019, Martínez-Rubio et al., 2025).

Depending on the cytometer, there are a number of markers that can be measured at the same time. Therefore, samples must go through this process several times to obtain information about the different IPT intensities. Preprocessing allows us to join and normalize all the data to obtain a matrix for each sample where the rows correspond to a cell and each column corresponds to each biomarker or cellular characteristic. Preprocessing of the data is necessary to finally obtain the merged data matrix. Other authors, such as Martínez-Rubio et al. (2025) or Chulián et al. (2023), have already preprocessed these data for other studies. We take this preprocessing as valid to work from it with our data. Furthermore, for each sample, we simplify the cells set to make the code faster using a random selection of cells, so all samples will have 10000 cells.

There is a peculiarity: In each bone marrow some biomarkers are not measured in others bone marrow, therefore the common basic markers must be found to distinguish ALL. We focus on samples which have the markers CD10, CD19, CD20, CD34, CD45 and SSC.A (Ortunõ Giner and Orfao, 2002, Martínez-Rubio et al., 2025). These biomarkers allow to distinguish our main subpopulations of cells along with maturation states in B lymphocytes, therefore, we eliminate from our study those who do not meet this requirement.

### 2.3 Treatment and outcomes

Currently in Spain, SEHOP-PETHEMA 2013 is the protocol used to evaluate patients diagnosed with leukemia. In that protocol, patients are identified in risk groups and they are treated depending on several values from their analyzed samples.

Treatment begins few days after the disease is detected and has different phases. Each phase corresponds to a set of drugs and blood and bone marrow samples extractions to monitor the progression of the disease in the patient. According to the protocol, a patient is considered to have responded correctly to treatment if there is less than 0.01% disease on day +15 and on day +33 no disease is detected. Data from day +15 are not available from hospitals in addition to there are very few cells in those samples. Otherwise, the patient is considered non-responsive to treatment and a change of risk group and other alternatives are studied to improve their therapy. Figure 2 shows a summary of the drug doses and patient follow-ups indicated in the protocol.

**Fig. 2.**
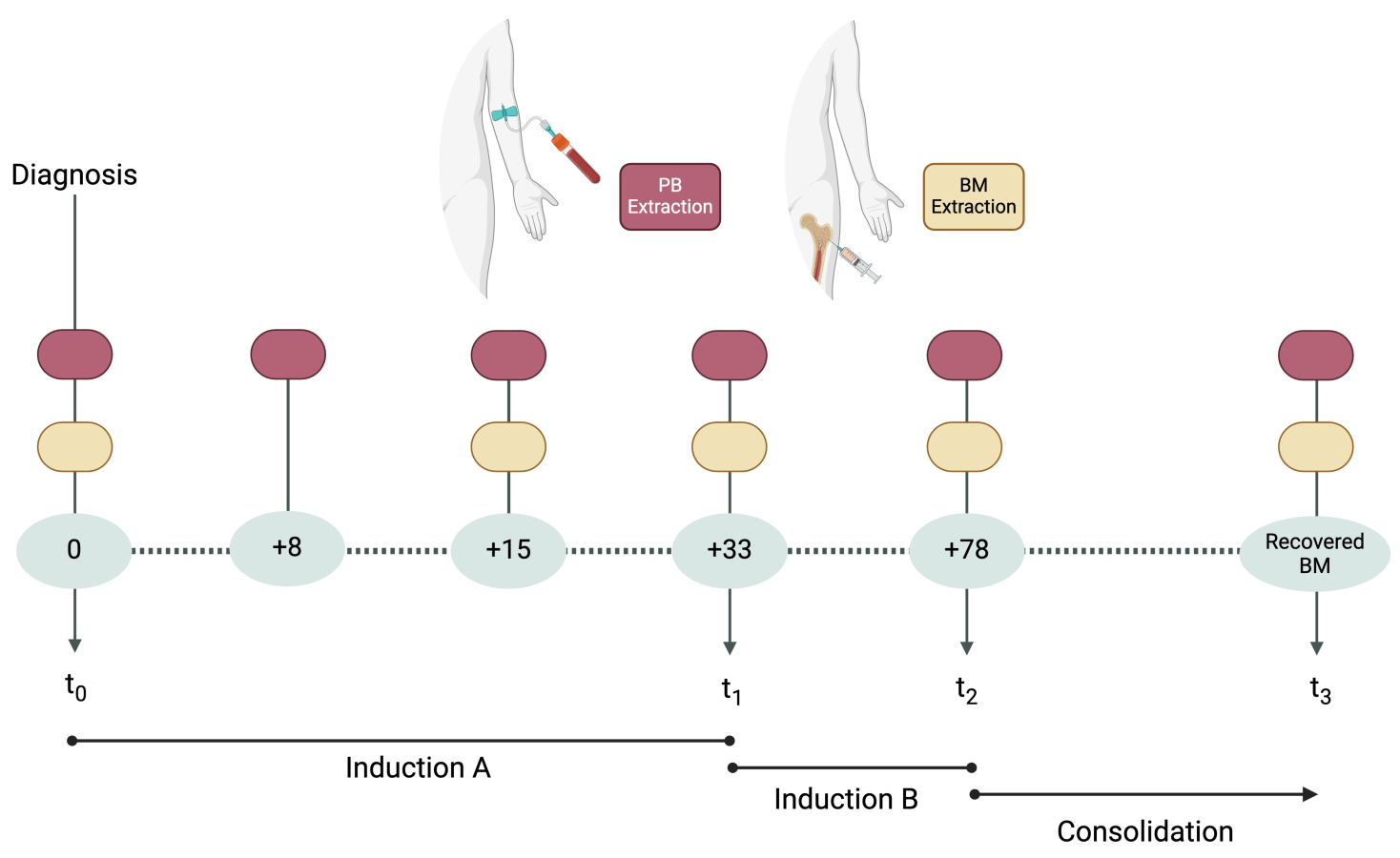
Chemotherapy schedule. Times in which bone marrow samples are obtained and studied. In this work, we use samples data from day 0, +33 and +78. It is possible to find some samples from recovered patients whose samples were extracted at least one year after treatment ended.

If the disease reappears once treatment has been established in a patient, the patient suffers a relapse. Over the years, there have been different protocols for the treatment of this disease, improving more and more but at present, there is a 20% relapse rate in patients diagnosed with B-ALL.

As indicated, we consider *t*_0_ as diagnosis day and *t*_1_ day +33 in addition to the day +78 of treatment associated to *t*_2_ (see Figure 2). On the other hand, if any patient who has recovered has had bone marrow extracted after finishing treatment, associated to *t*_3_ in Figure 2, that sample is taken as regenerated bone marrow. A recovered bone marrow must have the same behavior as a healthy bone marrow, therefore, it is possible to associate those samples with a normality in the bone marrow.

The aim is to analyze patterns by studying the data in bone marrow, which is where B-ALL originates. In this study we will consider the sample from the diagnosis of the disease, on day 33 from the start of treatment and the sample prior to beginning the consolidation phase of treatment, on day 78, approximately. We consider patients who respond to treatment, regardless of the fact of relapsing or not after the whole treatment is finished.

In addition to automating the cell selection process in the bone marrow, there are some medical questions related to identifying patients who relapse and those who do not relapse at the time of diagnosis (Martínez-Rubio et al., 2025, Chulián et al., 2023).

### 2.4 Exclusion criteria and study patients

From the 116 patients registered with a diagnosis of Acute Lymphoblastic Leukemia in the period of time covered by this study at the Niño Jesús Hospital in Madrid, we selected the patients who responded to treatment and for whom monitoring samples were available on days 33 and 78 of treatment. We added another restriction related to the minimum biomarkers that a sample must have to unify the way in which the different types of cells are identified, as it is indicated in Section 2.2. Finally, patients included in this study must have been diagnosed as Common B-ALL, in this way, we remove from the process those patients with other types of B Leukemia.

These criteria imply including 75 patients in the study of which 14 of them are relapses (R) and 61 non-relapses (NR). There are three time points for each patient included in the study along with the 9 samples considered regenerated bone marrow. Finally, 234 bone marrow samples are used in this work.

#### Limitations

There are several limitations related to patients’ data availability which must be acknowledged.

First, the retrospective nature of the study limits the standardization of sample collection and processing along with the study of the limited availability of flow cytometry files. Furthermore, follow-up data may be incomplete as a result of patient transfers to other hospitals, thereby reducing the reliability of longitudinal analyses. Additionally, the qualitative nature of clinical data extracted from medical reports hinders the definition of uniform subgroups, unlike the more structured cytometry data. Further, the limited range of available cytometry markers, discrepancies in cytometer settings, and batch effects restrict the depth of immunophenotypic characterization. To address this, we employed sample merging strategies (see Section 2.2).

Lastly, the dataset includes samples from 75 patients, with a marked imbalance between relapsing (R) and non-relapsing (NR) cases. This limited sample size, particularly in the relapse group, raises concerns about potential overfitting and limits the generalization of the findings. Due to the highly invasive procedure constituting bone marrow extraction, training of the regenerated patterns was only feasible using the 9 available samples from day +78, further emphasizing this constraint.

These limitations highlight the critical need to expand the patient cohort in future studies to ensure more robust, reliable, and generalizable results. Prospective validation efforts are currently underway and are expected to help address several of these limitations in future works.

### 2.5 Pipeline

We aim to classify cell subpopulations in patients’ samples using information from regenerated bone marrow cell samples. Data used in this study correspond to a set of patients’ samples for different times of the treatment. We consider to this end two pipelines (Figure 3):

**Fig. 3.**
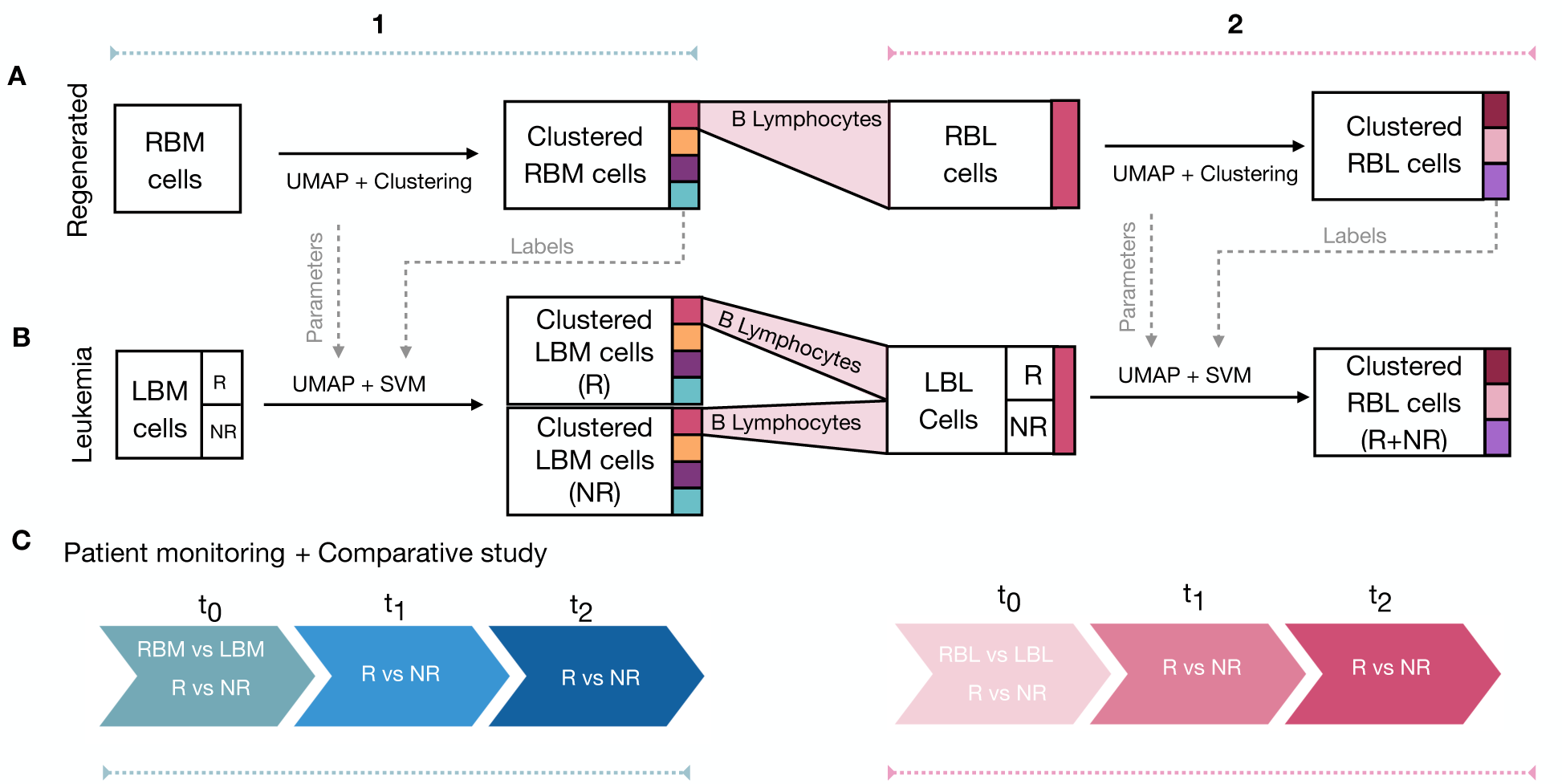
General pipeline. Steps conducted in the study. **A1**. Regenerated bone marrow (RBM) samples work as train dataset in the algorithm. UMAP and clustering techniques are applied to obtain transformed cells with their corresponding label. **A2**. B lymphocytes are selected (RBL) and UMAP is applied in their corresponding raw data to obtain clusters in this subpopulation. **B1**. UMAP is applied to leukemic (LBM) samples to obtain the dimensionality reduction associated to train dataset. Support Vector Machine (SVM) allows to cluster test dataset based on clusters in train dataset. **B2**. Selection of leukemic B lymphocytes (LBL) leads to analyzed these cells in the test dataset, using UMAP and SVM analogously, obtaining labeled B lymphocytes. **C1**. Bone marrow differences between relapses (R) and non-relapses (NR) at the three times along with cells comparison with RBM at diagnosis time. **C2.** LBL differences study between relapses (R) and non-relapses (NR). LBL at diagnosis compared with RBL.

– Pipeline A, which involves only regenerated bone marrow cells (RBM), Figure 3 A. These data are used as train dataset in our algorithm, where subpopulations in RBM are differenciated to select B lymphocytes (RBL) and study their maturation states cells subgroups.

– Pipeline B uses data from two groups of patients: relapses (R) and non-relapses (NR). For both two groups, leukemic bone marrow cells data (LBM) is used as test dataset (Figure 3B) based on the train dataset from Pipeline A (RBM). Analogously, data is processed to obtain leukemic B lymphocytes from the different groups, LBL (R) and LBL (NR). All leukemic cells are classified trough Support Vector Machine based on Pipeline A.

Furthermore, each pipeline leads to two parallel studies: Same processes are performed for the entire bone marrow (Figure 3.1) and for the B lymphocytes (Figure 3.2).

First of all, collected samples have been preprocessed as Martínez-Rubio et al. (2025) shows to obtain a single filtered file per sample. This process leads to obtain a dataset table where all the cells of all patients at all times are stored. We consider cells from regenerated bone marrow as train dataset while leukemic cells are considered as test.

As it is shown in Figure 3A1, cells in regenerated bone marrow samples are treated by UMAP to reduce their dimensions and then, DBSCAN clustering technique is applied to obtain four subgroups in that dataset. Therefore, there are clusters associated to cells which allow to select B lymphocytes (Figure 3A2) to obtain new clusters inside B cells.

To manage data from the rest of samples (Figure 3B1) UMAP is based on train dataset (RBM) and the clustering technique is based on zones from the train dataset clusters, therefore, we use Support Vector Machine (SVM) to obtain labels for those cells. Selecting B cells in the test dataset and applying UMAP and SVM (Figure 3B2) labels for maturation stages in B cells are obtained.

Finally, different analysis are presented to study (Figure 3C) both the behavior of the entire bone marrow (Figure 3C1) and development of the set of B cells in a regenerated setting and in leukemia (Figure 3C2) monitoring each patient.

After monitoring all the patients, we raise questions already asked by other authors such as Velasco et al. (2023), Fuster (2014) or Gong et al. (2024), who aim to find significant differences between patients who relapse and those who do not relapse at the time of diagnosis. We focused on carrying out the study of these groups of patients throughout the three time points available in our work, using basic statistical techniques, to establish a basis for more specific future study.

A pseudocode is included in the Supplementary Information to improve understanding of the code used to carry out all the steps described in the pipeline.

### 2.6 Dimensionality reduction and clustering techniques

Once the data has been filtered, the markers to study common to all samples are chosen: CD10, CD19 CD20, CD34, CD45, SSC.A. We seek to represent the different samples as simple as possible without losing information. For this objective, it is important to use dimensionality reduction techniques (Ujas et al., 2023, Wang et al., 2023) and some of them have been proved as a method to detect abnormal behavior of cells (Weijler et al., 2022, 2021) applied to myeloid leukemia.

Our work focuses on the UMAP technique since it provides fast and accurate method of data analysis, (Stolarek et al., 2022, Wang et al., 2021). This tool allows us to study the samples in a two-dimensional way, taking into account all the markers at the same time. Thus, the cells that are most similar will group together and those that form different groups will distance themselves depending on how different they are. The UMAP results allow to label each cell and obtain the initial information from it.

#### UMAP algorithm parameters

UMAP algorithm in *Python* allows to modify four basic parameters for its implementation. We highlight four hyperparameters which can significantly impact the result, see Table 2. These parameters are related to distance and neighbors in their corresponding subgroups.

**Table 2.** UMAP parameters values used for implementations.

| UMAP parameters | Meaning | Values |
| --- | --- | --- |
| n_neighbors | Cells neighbors | 50 |
| min_dist | Distance between cells | 0.01 |
| n_components | Dimensions | 2 |
| metric | Metric used to distances | Euclidean |

The metric used is Euclidean and we decide two dimensions to reduce the data since it could be visualized and managed in the plane. Those are values by default.

The goal is to find the distribution that most clearly distinguish these four cellular subgroups previously defined. Therefore, values have been selected depending on results from implementations with several parameters values mainly by modifying cells neighbors and the distance between cells. These parameter combinations performed can be found in Supplementary Information (SI).

Finally, we have considered values in Table 2 in this work.

#### DBSCAN cluster algorithm

DBSCAN is one of the most widely used clustering methods (Ye and Ho, 2019) and it is considered effective for working with large noisy databases (Ester et al., 1996).

To initialize the algorithm, two parameters must be defined: *ε* and *n*, corresponding to the maximum distance and the minimum number of neighbors, respectively. This method takes an observation and evaluates the number of points in its neighborhood whose distance is less than *ε*. If the number of neighboring events is greater than or equal to the defined parameter *m*, a cluster is formed. Additionally, each cluster has a central reference observation. Points that are not within a distance *ε* of any central observation are considered noise. Therefore, the effectiveness of this algorithm relies heavily on the proper definition of the parameters that determine the allowed distance *ε* and the minimum number of neighboring events *n* required to form a cluster.

Let *p* = (*p*_1_*, . . ., p_n_*) be a central observation and *q* = (*q*_1_*, . . ., q_n_*) a neighboring point within the same cluster. The Euclidean distance between them must satisfy:

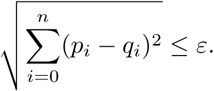

Depending on the parameters selected, we can obtain different number of clusters and amount of noise related a cells which are not identified with clusters. Once we have captured the UMAP data, we want it to be automatically clustered and associated with the four cellular subgroups previously defined (Figure 1). To find the optimal result, we perform iterations varying the parameters and use those that help to distinguish the subgroups of interest (see SI). In this case, values obtained as optimal for our study are *ε* = *eps* = 0.5, *m* = *min samples* = 25, using the definitions of *ε* = *eps* and *m* = *min samples* as shown in the code available.

#### Support Vector Machine classifier

Adjouadi et al. (2005) describes Support Vector Machine (SVM) as a powerful tool in the field of object classification and pattern recognition and in other works (Toedling et al., 2006, Rajwa et al., 2008), various machine learning tools tested for classification of scatter features of biological samples obtaining that SVM-based algorithms were especially promising. This method uses a training group of samples and it applies its algorithm to assign a label to new samples, obtaining groups of cells for the new sample associated to the training group. Throughout this study we used the default parameters.

The SVM algorithm finds the hyperplane that best separates the labeled classes in the feature space. Its goal is to maximize the margin, which is the distance between the hyperplane and the closest data points from each class, known as support vectors. By doing so, SVM aims to achieve an optimal separation that generalizes well to new, unseen data. As Cervantes et al. (2020) define, for the linearly separable case, *S* = [(*x*_1_*, y*_1_)*, …*(*x_l_, y_l_*)], if (*w, b*) is the solution to the following optimization problem:

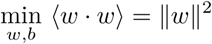

subject to *y_i_* (⟨*w* · *x_i_*⟩ + *b*) ≥ 1, then the maximal margin is given by 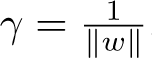.

In this case, the Radial Basis Function (RBF) kernel (one of the most widely used kernels in SVM applications) is applied. The kernel is defined as:

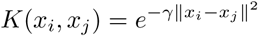

where the parameter *γ >* 0 controls the width of the Gaussian function and, consequently, the flexibility of the decision boundary.

This transformation allows the SVM to find a nonlinear decision boundary in the original input space while still solving a linear separation problem in the kernel-induced space. Furthermore, the selected method, *One vs One*, builds 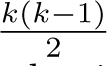 binary classifiers for a problem with *k* different classes. Each classifier is trained using only data from two classes at a time. When predicting the class of a new input *x*, all 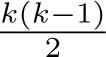 classifiers vote. Each classifier compares two classes and gives one vote to the predicted class. For example, if a classifier trained to distinguish class *i* from class *j* says that *x* belongs to class *i*, then class *i* receives one vote. The final predicted class is the one that obtains the highest number of votes. A main drawback of the *One versus One* method is that if the input lies in an ambiguous region (i.e., near decision boundaries), it might receive the same number of votes for two or more classes, making the final decision uncertain (Cervantes et al., 2020).

In this study, we consider 9 regenerated bone marrow samples that serve as training. From the classification obtaining for those regenerated bone marrow, SVM application allows to identify subgroups in the rest of samples. According to the clustering obtaining previously for those cells, it is possible to automate the subgroup selection process.

#### Data analysis techniques

It should be noted that to apply a test of significant differences between two groups, it is necessary to check whether the samples follow a normal distribution. In this case, some samples have a non-normal distribution, therefore, we apply *Mann-Whitney U* (MWU) test, a non-parametric version of *T-Student* test.

The MWU test, also known as the *Wilcoxon rank sum* test, is used to compare differences between two groups on an ordinal variable without a specific distribution. It is a nonparametric test, whereas the independent samples *t-test* is parametric and requires the variable to be measured at the interval or ratio level, in addition to being normally distributed (Nachar, 2008). Both tests assess whether two groups differ on a continuous variable, but they differ in their distribution assumptions. The MWU is suitable when the parametric assumptions of the t-test are not met (Ruxton, 2006). This test is recommended when the variances between groups are not equal, showing better control of Type I and Type II errors. It is argued that the unequal variance t-test is more reliable and should be preferred over conducting preliminary homogeneity of variance tests, which can decrease statistical accuracy. Additionally, it mentions its implementation in common statistical software and its application even when the underlying distributions are not normal. This test, also known as the Welch test, is recommended as the most robust and practical option in most scenarios, outperforming alternatives in terms of error control and ease of use.

The Bootstrap method is a statistical technique used to assess the uncertainty of estimators in complex situations, complicated data, or small samples where standard models may not be applicable. It relies on resampling with replacement to estimate the distribution of a statistic and derive confidence intervals. By generating multiple random samples from the original data, it calculates the sampling distribution of the statistic of interest, allowing the evaluation of properties such as bias and variance. The main bootstrap confidence intervals include the percentile, studentized, and BCa (bias-corrected and accelerated) intervals, with the latter improving accuracy in the presence of skewed distributions (Davison and Hinkley, 1997).

Bootstrap is applied in various fields, such as regression and sample comparison, providing alternatives to traditional parametric approaches. Techniques have also been developed to improve variance estimation, such as the double bootstrap and delta method. Additionally, bootstrap is classified into parametric, which assumes a specific distribution, and non-parametric, which does not rely on distributional assumptions. The advantages of bootstrap include its simplicity and applicability to complex estimators, often providing more accurate confidence intervals than traditional methods based on normality assumptions. However, its effectiveness depends on the estimator used and the problem conditions. Finally, comparative studies have shown that advanced methods like BCa achieve second-order accuracy, improving estimation in finite samples.

Finally, in order to manage the data resulting from applying the different techniques mentioned above, a study of differences between different sets of data is proposed.

### 2.7 Data and code availibility

The source code and functions used in this article can be consulted at https://github.com/ananinolopez/SI NinoLopez25. This repository also includes the full tables from the study of the patients and other figures related to iterations in the code. Additionally, pseudocode is available in this Supplementary Information.

The code was executed on an iMac with the following specifications: Retina 5K, 27-inch (2020), equipped with a 3.3 GHz 6-core Intel Core i5 processor, AMD Radeon Pro 5300 graphics with 4 GB VRAM, 64 GB of 2667 MHz DDR4 memory, running macOS Sequoia 15.5. Execution time for the completed code was 2 hours and 34 minutes. The algorithm is proposed as a tool that, in clinical settings, would process the code in just two minutes per sample.

## 3 Results

The results exposed are based on two parallel studies: Both processes are performed for the whole bone marrow dataset and for the B lymphocytes cell group. Furthermore, each result is related to the step shown in Figure 3.

### 3.1 UMAP and DBSCAN analysis provides an automatic cell identification for regenerated bone marrow subpopulations and B lymphocytes maturation states

We aim to classify regenerated bone marrow (RBM) and regenerated B lymphocytes (RBL) automatically, which we previously defined as cells from patients who ended treatment and recovered the normal cell levels.

Firstly, using the Pipeline A1 in Figure 3, we have used the UMAP dimensionality reduction algorithm to obtain a minimal cell data representation in RBM samples. This allows us to distinguish four main groups related to their immunophenotypic signature (erythroblasts, myeloid cells, B lymphocytes, and monocytes and T cells, see Figure 1). This signature can be interpreted as the immunophenotypic marker intensities in each subgroup, which leads to the cell subpopulation patterns indicated in Figure 4A. Cells that correspond to the myeloid lineage have a higher CD66 intensity in comparison to the remaining RBM cells. A higher intensity of CD19 fluorescence relates to B lymphocytes, and those with high CD45 but low CD19 are associated to monocytes and T cells. Finally, we can verify that the erythroblasts dataset are also in agreement with the previous biological definitions in terms of fluorescence.

**Fig. 4.**
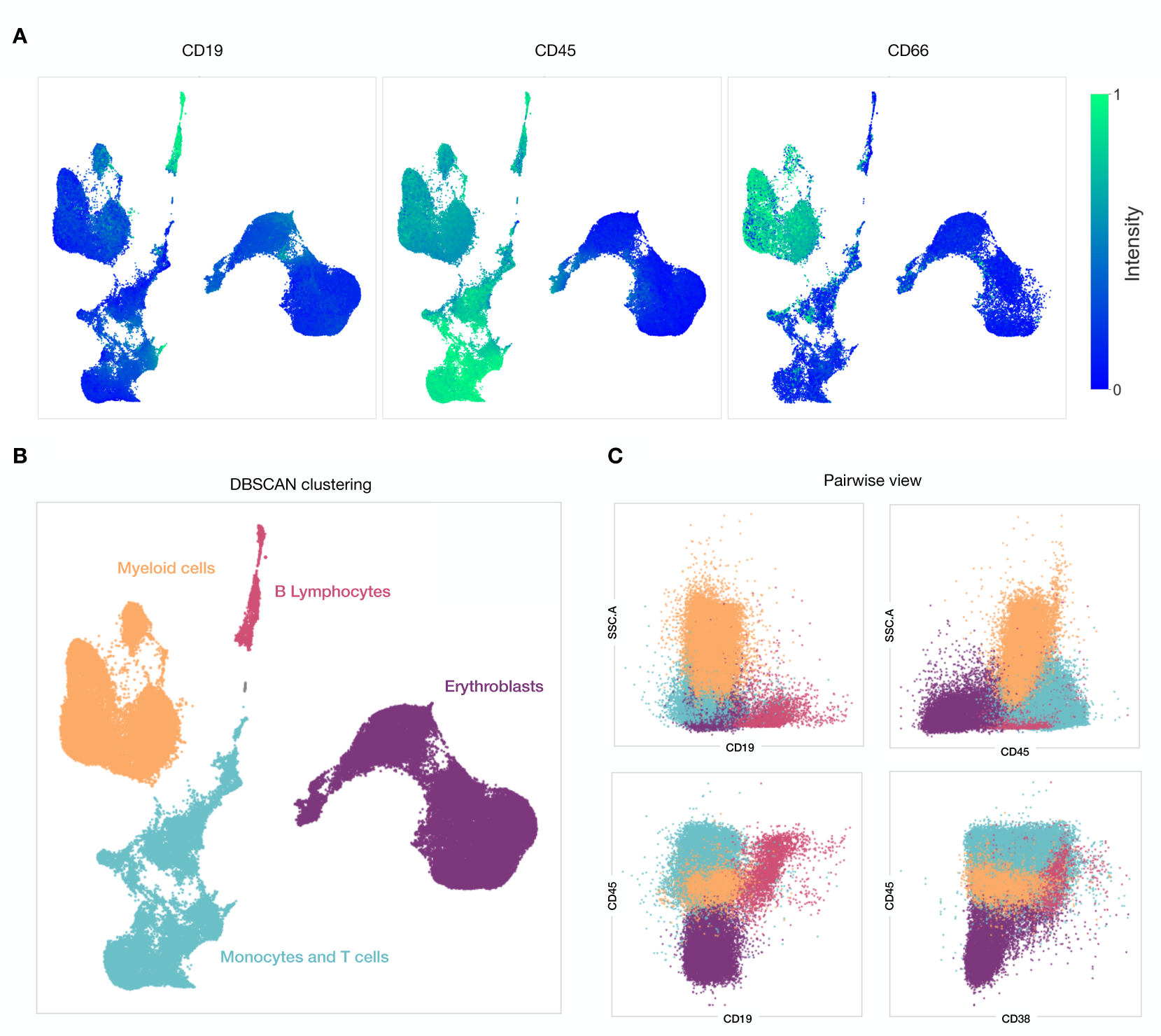
Automatic classification of regenerated bone marrow (RBM) subpopulations through UMAP and DBSCAN algorithm. For this result, we use the steps related to Pipeline A1 in Figure 3. **A.** In the UMAP representation of the RBM data we can highlight the intensity of each original IPT marker to find cells with different characteristics. **B.** DBSCAN algorithm is used to cluster the data subpopulations, which are labeled depending on the IPT intensities in each subgroup. **C.** Using the information from the obtained clusters, we represent the original IPT markers space pairwise. The cell subpopulation arrangement agrees with the bone marrow two-dimensional visualization used in routine clinical analysis.

Later on, the DBSCAN technique has been applied to cluster the previous data obtained from the UMAP reduction. As observed in Figure 4B, we can automatically obtain four groups related to the four main cellular subpopulations in RBM cells. Now we aim to check that the defined subpopulations agree with the biological definitions of the immunophenotypic signature of the different RBM cells (Orfao et al., 2019). For this purpose, we display the same data two-dimensionally (see Figure 4C), now using the original intensity fluorescence markers. We verify that the UMAP-labeled subpopulations correspond to either erythroblasts, myeloid lineage cells, B lymphocytes, or monocytes and T cells.

For the case of B cells, the clusters previously obtained from Pipeline A1 allows us to select in Pipeline A2 the RBL cells in study (Figure 3). Once B lymphocytes are labeled (see Figure 5A), we have used the original IPT markers and reduce the information to a two-dimensional space applying UMAP again, now only with the RBL cells. A representation of IPT intensity in the UMAP reduction of RBL cells space is observed in Figure 5B.

**Fig. 5.**
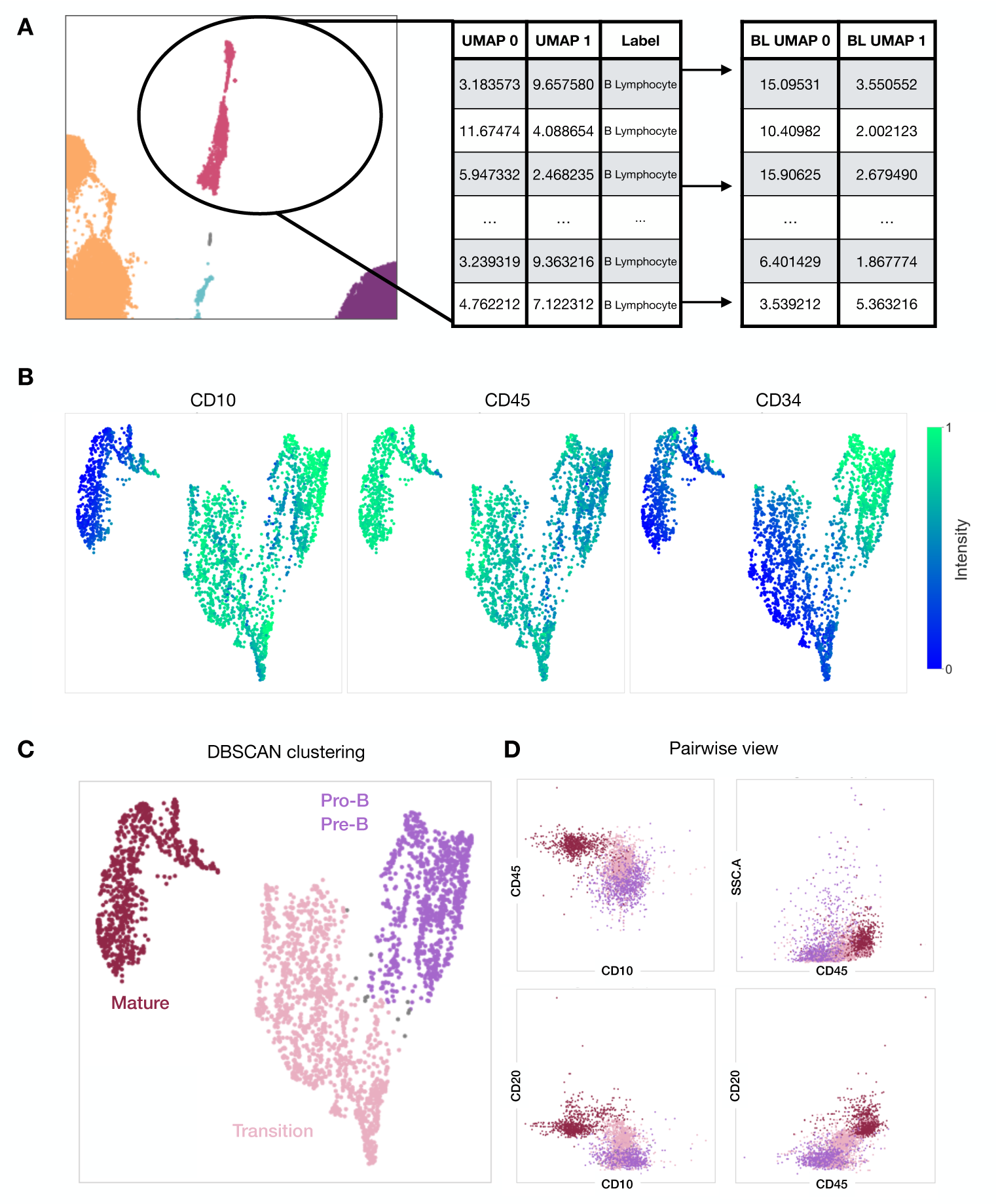
Study of regenerated B lymphocytes (RBL). Steps related to Pipeline A2 in Figure 3. **A.** Cells labeled as B lymphocytes are selected and UMAP is implemented again. **B.** IPT intensity in each cell. As the cell develops, it acquires CD45 while losing the intensity of CD10 and CD34. **B.** DBSCAN is applied to the set of cells and three subpopulations can be distinguished. These correspond to maturation stages. **D.** Bidimensionally view of data checks the differentation obtained.

We can observe three different groups, which correlate to the the maturation states presented in Figure 5C. A first group, with relatively high CD45, and very high CD10 and CD34 intensity, corresponds to Pro-B and Pre-B cells. A second group with a very high CD45, low CD34 and high CD10, is associated with Transition RBL cells. The last group, are those cells with very high CD45, now with low CD34 and CD10, compatible with Mature RBL cells. These three clusters are the ones obtained in the DBSCAN algorithm, and they are associated to the different maturation states in RBL subpopulations. In order to verify the obtained cell classification, the original IPT markers of the cells are represented two-dimensionally in Figure 5D. Thus we check that cells are distributed as the biological definitions of the RBL cells maturation states Orfao et al. (2019).

We are aware that the use of other techniques such as PCA, t-SNE or UMAP can be considered for this type of analysis. However, as seen in Park et al. (2025), the authors performed a comparison of both UMAP and t-SNE techniques for flow cytometry data, for the early detection of MRD in leukemias and lymphomas. Both methods resulted in a 95% agreement with the manual gating, sensitivity, and negative predictive value. Nevertheless, we have performed an additional comparison between UMAP and PCA, and the use of both k-means and DBSCAN (see Figure 6). After this analysis, either with k-means or PCA, the bone marrow cell subpopulations are not clearly identified. Therefore, due to the ability of UMAP to keep the global and local structure of the dataset, we keep both UMAP and DBSCAN in the following results to maintain the biological reasoning.

**Fig. 6.**
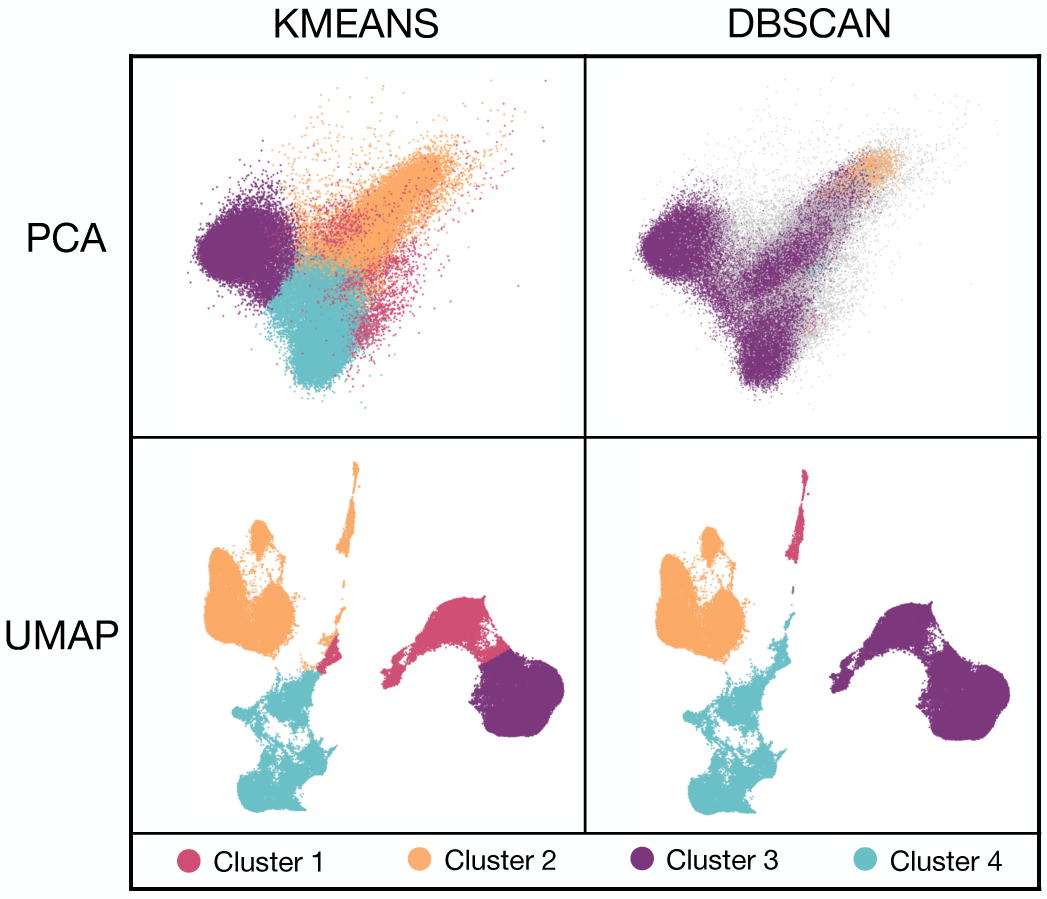
Comparison of dimensionality reduction and clustering methods. **PCA + K-means**: Three main cell populations are grouped, although the distinction between them is not clearly defined. Additionally, there is a group of cells not distinguishable in the projection. **PCA + DBSCAN:** This approach distinguishes only two cell populations. DBSCAN identifies most cells as erythroblasts, indicating a loss of resolution in separating cell subtypes. **UMAP + K-means:** UMAP provides a clearer definition of the cellular subpopulations in the two-dimensional space. However, when K-means is applied with *k* = 4, the resulting clusters do not correspond well with the four visually distinct subpopulations. **UMAP + DBSCAN:** This combination proves to be the most effective. UMAP clearly separates the four cellular subpopulations, and DBSCAN accurately identifies the four groups formed, aligning well with the visual distribution.

### 3.2 Algorithm for the identification of cell subpopulations

Once the RBM and RBL cells’ distribution is obtained, the goal is to associate each leukemic cell to a cellular subgroup, as indicated in Pipeline B in Figure 3. From the training dataset, this is, the RBM and RBL cells UMAP distribution, we obtain labeled zones (see Figure 7) through the Support Vector Machine (SVM) algorithm. Once this is performed, we reduce the leukemic cell data (LBM and LBL cells) to the prior UMAP coordinates and label each leukemic cell according to the previously defined zones.

**Fig. 7.**
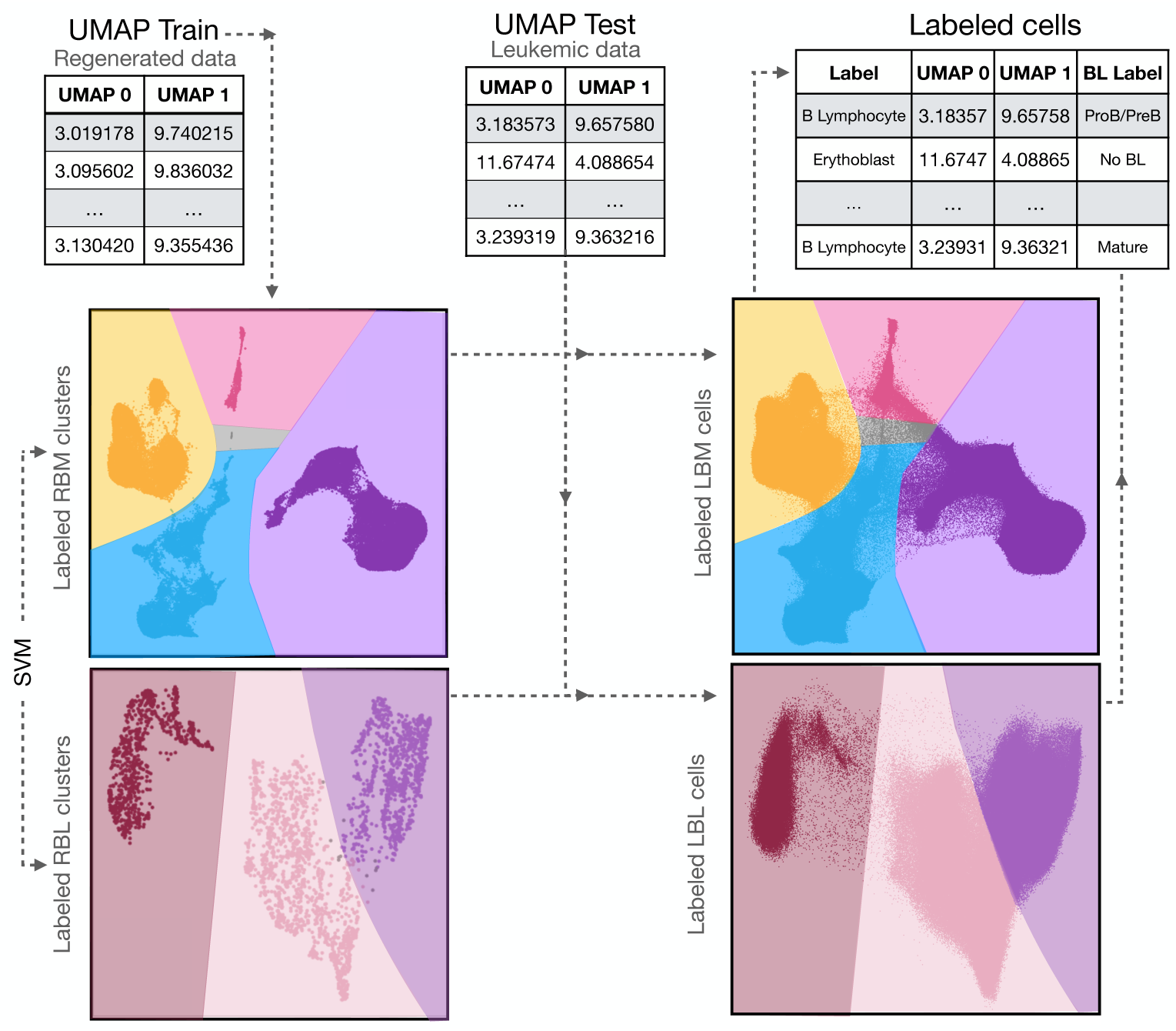
Automatic labeling of leukemic cells by SVM. The steps followed are those related to Pipeline B, Figure 3B. The UMAP distribution of RBM cells can be trained to obtain hyperplanes using SVM, which can distinguish several cell zones or clusters. Using the UMAP coordinates from the RBM cells, LBM cells dataset can be reduced, and later be used as the test dataset. With the clusters previously obtained, labels are derived for the LBM cells. Analogously, LBL cells can be then labeled by applying the same process to RBL cells.

Given the absence of true cellular labels for supervised validation for both RBM and LBM cells, we assessed the performance of the SVM classifier on the training set of RBM cells. The resulting confusion matrix shows that the method correctly classifies the RBM cell subpopulations, indicating that the model has completely learned to distinguish the groups previously defined by UMAP and DBSCAN, based on RBM cells. This outcome suggests that the clusters are separable in the transformed feature space, supporting the validity of the initial segmentation. Nevertheless, as the evaluation was performed on the training data, the possibility of overfitting cannot be excluded.

### 3.3 Bone marrow composition monitoring over time

As indicated in Section 2, three time points have been collected from each patient included in this work, directly leading to patient monitoring. We can then analyze the patients’ samples over time and compare them with the RBM distribution, which is stable over time as it represents the normal cell levels in a recovered patient. For instance, in Figure 8A, three time points from a single patient are represented and compared with a RBM sample. Considering all RBM samples and all LBM samples, we can compare the subpopulation percentages from the RBM cell distribution with the evolution of all patients at each time point. As shown in Figure 8B, we can thus analyze how the LBM cells generally behave at different times.

**Fig. 8.**
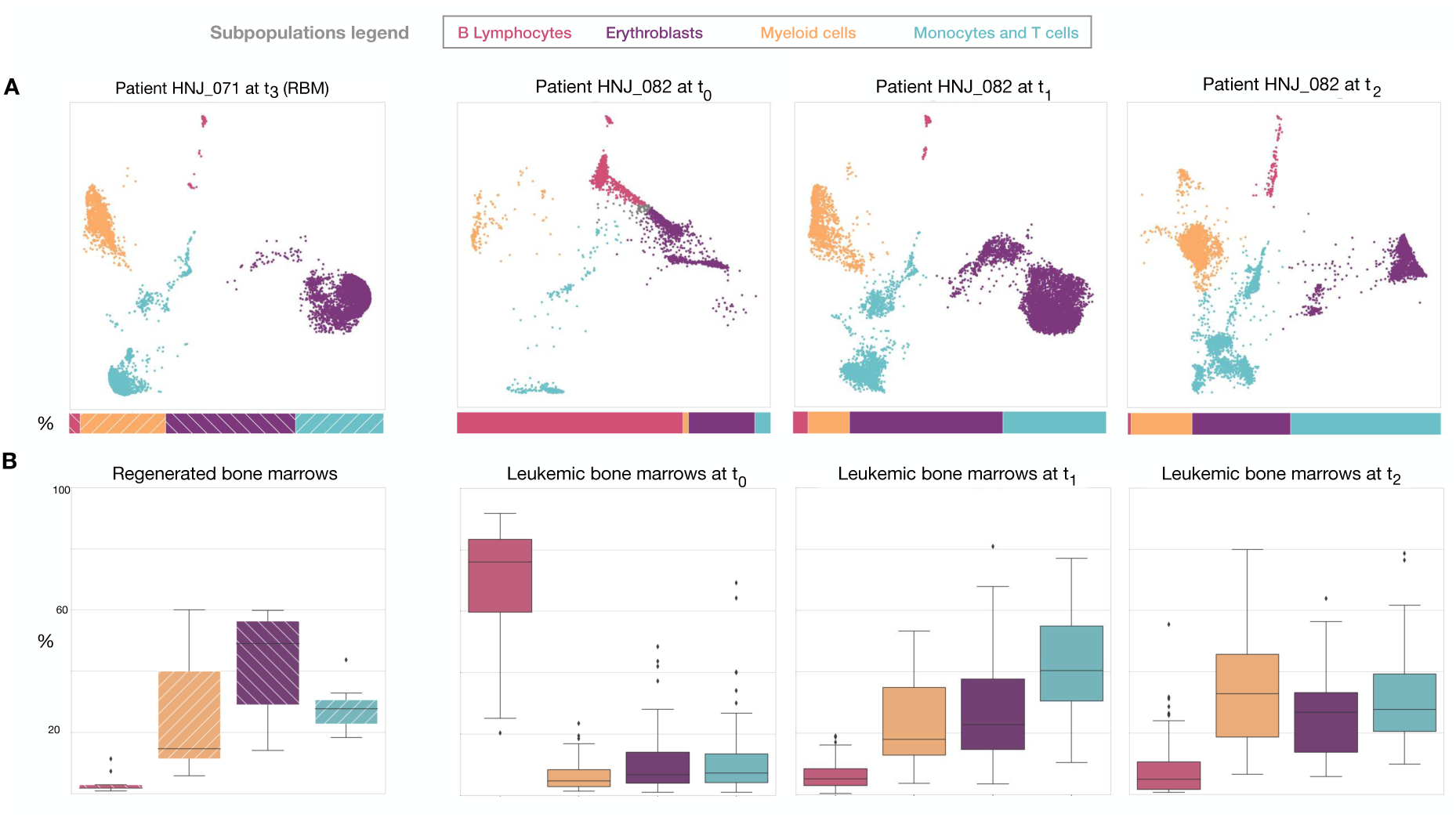
Comparison between RBM and LBM samples over time. **A**. For instance, we consider a RBM sample (HNJ 071) and compare it with samples from a patient (HNJ 082) over their three time points (*t*_0_*, t*_1_*, t*_2_). **B**. The composition of all RBM samples and all LBM samples at each time point are represented using box and whisker plots (stripped and plained boxes, respectively).

In Figure 8B, it can be observed that LBM samples at time *t*_0_ have an abnormal amount of B lymphocytes in comparison to RBM samples. This result is consistent with the classical behavior of the disease, due to the bone marrow being invaded by the malignant cells. We have also perform this analyzes for the RBL versus the LBL cells. We verify that LBL cells is mostly composed of Pro B and Pre B cells than mature ones. This aligns with the idea that in B-ALL, the cells that invade the bone marrow are immature B lymphocytes. Full data with the percentages of each subpopulation for each patient sample related to Figure 8 are included in Tables S1-S7 in Supplementary Information (SI).

### 3.4 Recovery processes for relapsed versus non-relapsed ALL patients

We have labeled the data considering the relapse or non-relapse status of ALL patients after therapy. The data consists of each patient’s bone marrow distribution over three different time points before the possible relapsing event. Our aim is to find differences over time between relapsed (R) and non-relapsed (NR) patients in their recovery processes. We could then analyze the LBM and LBL samples distribution (Figure 3C).

Using the RBM and LBM cells, we have applied Pipeline A and Pipeline B (Figure 3A and B) to both cohorts of patients, thus automatically labeling over time LBM and LBL cells for R and NR patients. We propose two different studies about the bone marrow datasets: In the first one, we have calculated and compared the percentages of each bone marrow subpopulation between the different cohorts of patients. On the other hand, from each IPT marker in each sample, we calculated the mean fluorescence intensity (MFI) corresponding to each cellular subgroup. Therefore, we have mean values related to each IPT marker for each subpopulation for each patient sample. With both pieces of information, we can perform a test to detect whether there are significant differences between both groups of patients. These strategies have been also used to analyze cells from patients on treatment days 0, +33 and +78 (*t*_0_*, t*_1_ and *t*_2_, respectively).

In the first study, we obtained for both cohorts the percentages of each LBM subgroups (see Tables S1-S7, SI) and LBL states (see Tables S8-S14, SI) at each treatment time point (*t*_0_*, t*_1_ and *t*_2_). With such information, we studied whether there are significant differences between R and NR patients and also between them and RBM, using *Mann-Whitney U* (MWU) test, thus deriving a *p-value* for each cellular subgroup, for each time *t*_0_*, t*_1_ and *t*_2_.

For the case of LBM cells (specifically for B lymphocytes and erythroblasts in *t*_2_) *p-values* with less than 0.05 were found (see Table S15, SI). We then calculate each 95% confidence interval for each group and show all results in Figure 9 related to Bone Marrow composition percentages. We highlight specifically that, for LBM cells at *t*_2_, the percentage of B lymphocytes in R patients (orange) approximates to the range of RBM cells, while in the case of NR patients (blue), their confidence interval is higher in mean.

**Fig. 9.**
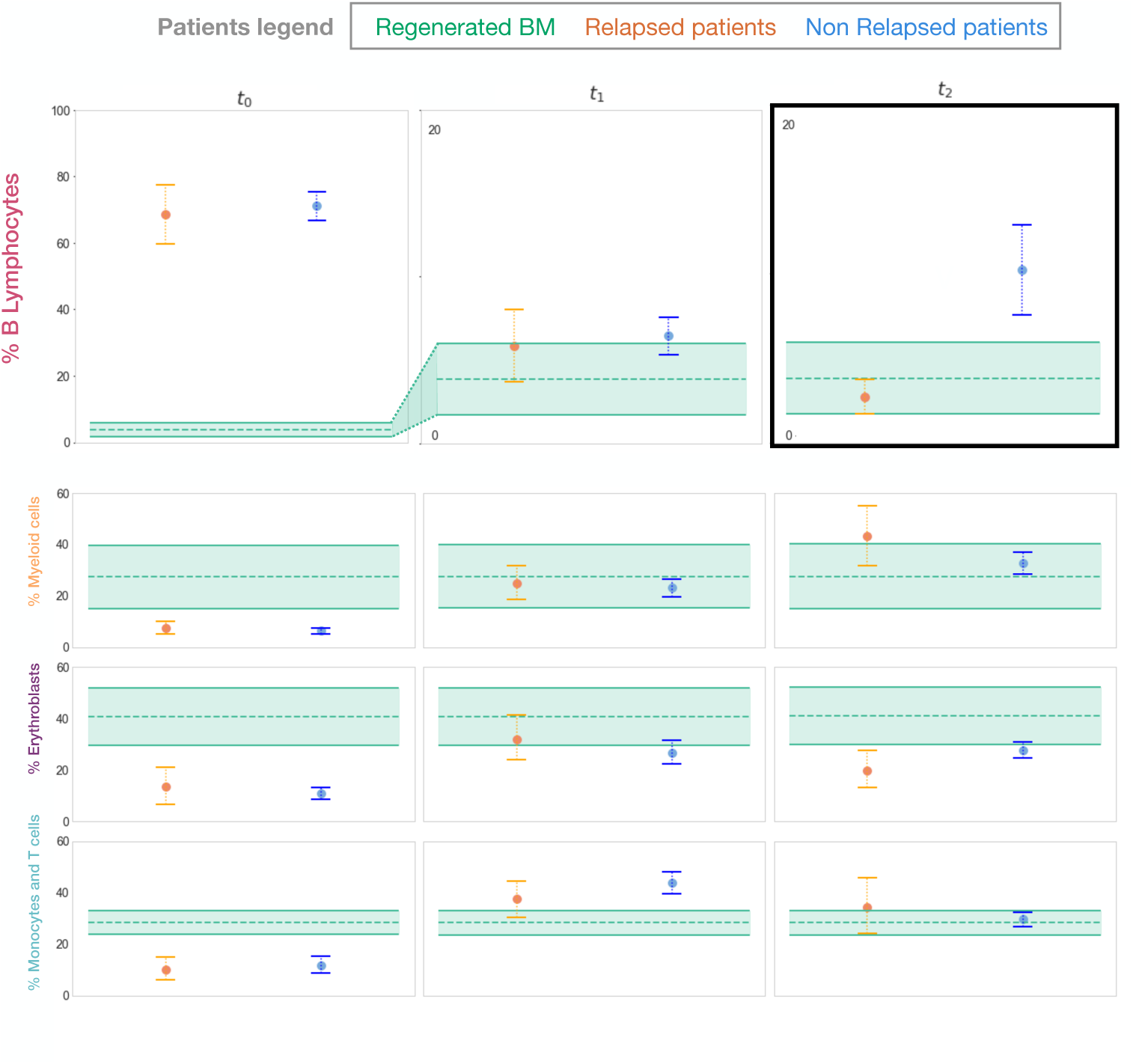
Confidence Intervals of each group of patients according to their LBM subpopulation proportions over time. A graphical representation of the 95% confidence intervals has been made for each time point and LBM subpopulations (B lymphocytes, Myeloid cells, Erythroblasts and finally Monocytes and T cells) for each group of patients: relapsed (orange), non-relapsed (blue) and regenerated (green band) patients. The mean value for relapsed and non-relapsed patients is represented by a dot while the RBM mean is represented by a dashed line. In all three cases, the solid horizontal lines represent the interval range.

We repeated this process with the percentages of the maturation states of LBL cells, obtaining some differences between R and NR patients in the UMW test, as well as their respective differences with the RBL cells. On the one hand, higher states (Transition and Mature) at *t*_2_ show differences between R and NR groups. Conversely, NR patients LBL cells compared with RBL presented most *p-values* are less than 0.05, while R patients present no differences at *t*_1_ and *t*_2_ in comparison to RBL cells. All these tests can be found in SI (see Tables S16-S17, SI).

In the second study, the same process was performed for both cohorts, but now deriving the MFI values of the IPT markers at *t*_0_*, t*_1_ and *t*_2_, both for each of the LBM subgroups (see Tables S18-S29, SI), as well as the LBL maturation states (see Tables S30-S38, SI).

According to the previous results for the percentage of LBM subpopulations, we have focused on the B lymphocytes and erythroblasts MFI (see Tables S39-S42, SI), since they are the subpopulations with more significant differences between both groups of patients. While we obtained no relevant differences between patients in the MFI values of the erythroblasts, we found some relevant differences in the MFI from the B lymphocytes subpopulation, specifically at the moment *t*_2_. There are significant differences between both cohorts of patients, according to the obtained *p-value* related to CD19 MFI (see Table S39, SI). On the other side, considering the differences between each group concerning RBM cells, there are significant differences according to CD19 and CD20 MFIs between NR patients and RBM, while there are no differences between R patients and RBM, as shown in Figure 10, (see Table S40, SI).

**Fig. 10.**
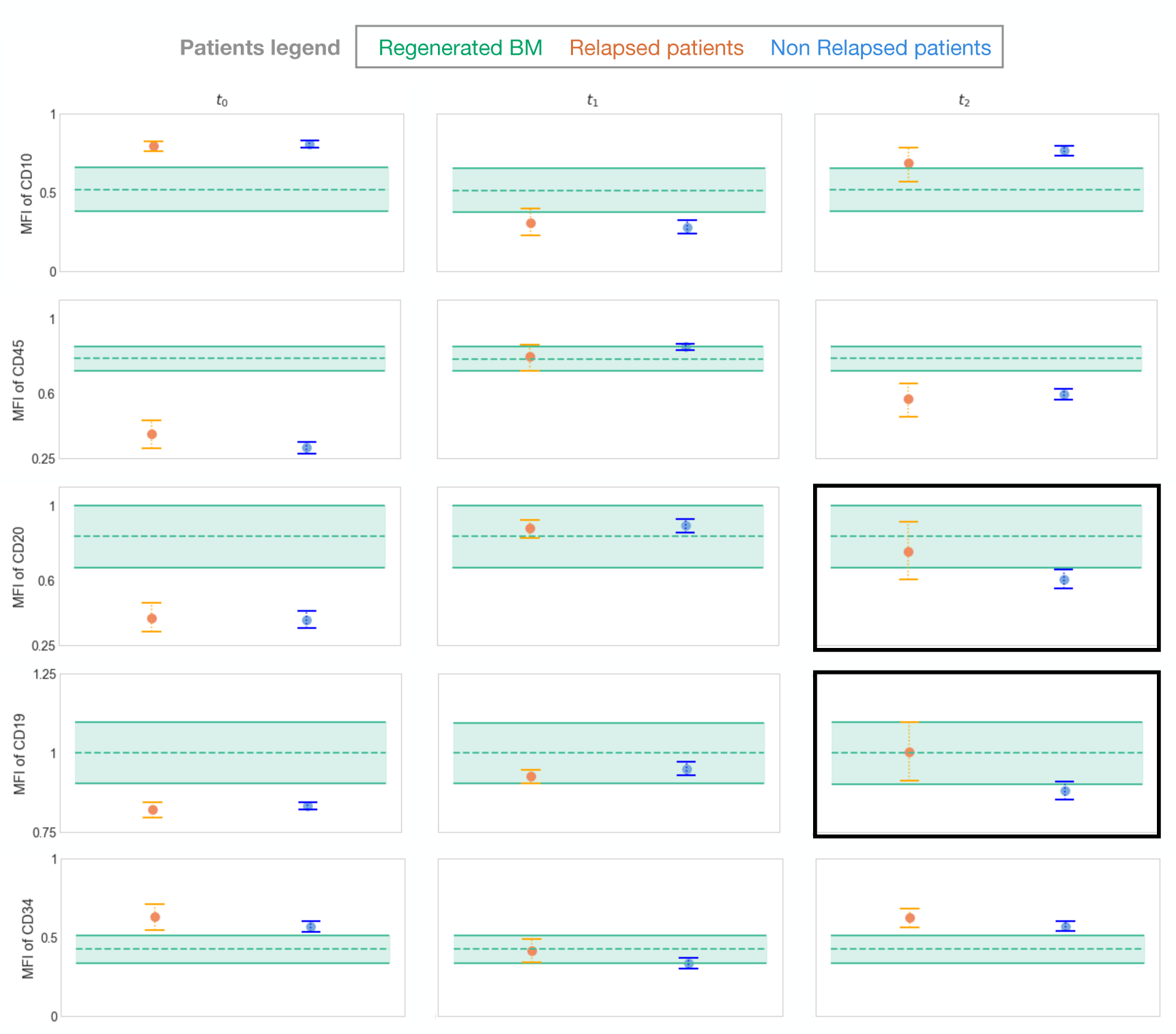
Confidence Intervals of each group of patients according to the MFI in their B lymphocytes. A graphical representation of the 95% confidence intervals has been made for each time point and bone marrow subpopulation for each group of patients: relapsed (orange), non-relapsed (blue) and regenerated (green band) patients. The mean value for relapsed and non-relapsed patients is represented by a dot, while the RBM mean is represented by a dashed line. In all three cases, the solid horizontal lines represent the ends of the intervals. MFI values are normalized between 0 and 1 and they are associated only to B lymphocytes in LBM.

Analogously, now repeating the process for LBL, our algorithm is applied to the study of the differences between R and NR patients’ B lymphocyte maturation states MFI. As previously stated, significant differences in the percentage of LBL maturation states were found only at the moment of *t*_2_. In this case, pro-B/pre-B and mature cells CD19 MFI, as well as transition cells CD34 MFI, reveal significant differences between R and NR patients at the moment *t*_2_ (see Table S19, SI). Specifically in CD19 MFI for pro-B/pre-B and CD34 for transition cells, R patients show significant differences between LBL and RBL cells, while NR patients present marginally significant differences in comparison to the CD19 MFI from RBL mature cells (see Tables 43-48, SI).

### 3.5 Classification approach

Based on the result below, we propose a simple classifier to assign a label to patients using the MFI values of CD19 and CD20 for B lymphocytes. The idea behind this classifier is the most intuitive one that arises from the observation of the previous result (Figure 10). For any clinician, this result will be of interest if the predictive power of the described intervals as classifiers is validated. Therefore, we propose distinguishing patients based on the MFI of each biomarker.

We take as reference the values from Tables S22 and S26 (in the Supplementary Information). From the MFI values of CD19 and CD20 at *t*_2_, we will compute the 10th percentile of the R group (*P*_10_*_R_*) and the 90th percentile of the NR group (*P*_90_*_NR_*). Using these values, along with their mean, we will study the classification of patients by proposing three different threshold values:

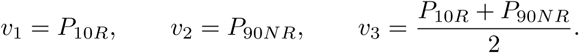

For each threshold value *v_i_*, with *i* = 1, 2, 3, the label assigned to a patient will be R if their value is higher than *v_i_* and NR otherwise. We now present in Table 3 the results of the classifier using our same patient cohorts. We include as well the confusion matrix and the accuracy to summarize the information for better comparison.

**Table 3.** Confusion matrix and accuracy from each classifier. Variation of *v*_1_, *v*_2_, and *v*_3_ for biomarkers CD19 and CD20 to compare their accuracy.

| Marker | Threshold | Confusion Matrix | Accuracy |
| --- | --- | --- | --- |
| CD19 | $v_1 = 0.809$ | $\begin{bmatrix} 12 & 2 \\ 48 & 13 \end{bmatrix}$ | 0.385 |
| | $v_2 = 1.04$ | $\begin{bmatrix} 6 & 8 \\ 7 & 54 \end{bmatrix}$ | 0.750 |
| | $v_3 = 0.925$ | $\begin{bmatrix} 8 & 6 \\ 14 & 47 \end{bmatrix}$ | 0.687 |
| CD20 | $v_1 = 0.441$ | $\begin{bmatrix} 12 & 2 \\ 48 & 13 \end{bmatrix}$ | 0.385 |
| | $v_2 = 0.89$ | $\begin{bmatrix} 4 & 10 \\ 7 & 54 \end{bmatrix}$ | 0.725 |
| | $v_3 = 0.666$ | $\begin{bmatrix} 6 & 8 \\ 18 & 43 \end{bmatrix}$ | 0.610 |

Given the variety in the accuracy results from Table 3, we generated a Random Forest classifier which includes the information from both biomarkers. However, due to the very limited amount of data, the model does not learn sufficiently. Given the constraints of the sample size, we applied a stratified 5-fold cross-validation to avoid selection bias. Additionally, we used oversampling to improve the training data by duplicating patient samples. Since the results remain similar, this suggests that more diverse data are needed, as oversampling only replicates existing samples. This issue is noted as a limitation and remains as future work. The metrics obtained for the Random Forest classifier, both without and with oversampling, are shown in Table 4.

**Table 4.** Random Forest classifier under different conditions. Random Forest was applied without oversampling, resulting in an accuracy of 0.733. Oversampling was then used to balance the dataset, but the accuracy remained the same when using a standard train-test split. Finally, we applied cross-validation with oversampling, which yielded a higher accuracy of 0.76.

| Method | Confusion Matrix | Accuracy |
| --- | --- | --- |
| Without oversampling (Train/Test) | $\begin{bmatrix} 1 & 2 \\ 2 & 10 \end{bmatrix}$ | 0.733 |
| With oversampling (Train/Test) | $\begin{bmatrix} 1 & 2 \\ 2 & 10 \end{bmatrix}$ | 0.733 |
| With oversampling (Cross-validation) | $\begin{bmatrix} 5 & 9 \\ 9 & 52 \end{bmatrix}$ | 0.760 |

## 4 Conclusions

Flow cytometry has been incorporated into clinical practice for decades for the diagnosis and monitoring of patients with hematological diseases. It has become a powerful and routine tool for sensitively identifying and quantifying the presence of pathological cell populations. Today, it is the technique of choice in many protocols for quantifying levels of measurable residual disease (MRD) during treatment. The quantification of MRD by flow cytometry has allowed tailoring therapies to patients, contributing to the improvement of the results of different protocols in recent decades.

Patients selected in this study present common characteristics related to available samples, B-ALL subgroup, treatment applied along with flow cytometry data from their samples. We have focused our work on the mathematical analysis of such flow cytometry data at different time points, specifically at the moment of diagnosis and when MRD quatification was performed, according to the SEHOP-PETHEMA protocol. Then, preprocessing allows us to normalize data and manage 234 samples from 75 selected patients. In these samples, regenerated bone marrow (RBM) and leukemic bone marrows (LBM) from different patients were included, this last with their label as relapsed and non-relapsed patients. Following the structured process presented in the general pipeline (Figure 3), the study of patients has consequently followed two main lines. In the first one, an algorithm is presented to classify cells from patients who have finished their treatment and they have recovered normal levels in their bone marrow. Using this classification, regenerated B lymphocytes are selected (RBL cells) and the same process was applied to distinguish maturation states in B cells. On the other hand, an analysis is presented about data from leukemic cells based on the regenerated bone marrow process.

As a first line of results, we obtained a two-dimensional representation of RBM cells by UMAP dimensions reduction (Figure 4). B lymphocytes, myeloid cells, erythroblasts and monocytes and T cells were reflected in the UMAP subgroups. To obtain defined subpopulations in bone marrow automatically, DBSCAN clustering technique was applied. The four groups related to such subpopulations in regenerated bone marrow allow us to study the bone marrow flow cytometry data from another perspective. As such, we have represented all cell data in two dimensions, while in clinical routine this process is usually performed by analyzing pairwise biomarkers. This would improve the technique of identifying subpopulations thanks to the reduction of dimensionality without losing the information of the cells analyzed. From these four groups, we selected the group of B lymphocytes and we analyzed the information about their cells using the previous process with UMAP and DBSCAN. Analogously, automatic classification related to B lymphocytes (RBL) allows us to identify cells’ maturation states (Figure 5). It can be observed that the distribution of B cells aligns with the immunophenotypic intensities of each maturation state. All pairwise representations of such immunophenotypic markers correspond with the gating techniques used in flow cytometry for the diagnosis and monitoring of ALL patients.

Once both RBM and RBL cells were labeled, we obtained a projection of the bone marrow which allows us to reproduce the labeling for leukemic cells (Figure 7). This algorithm uses data from RBM cells UMAP to reduce leukemic cells embedding data to two dimensions, and then, it labels these leukemic bone marrow cells (LBM cells) depending on their location in the UMAP by Support Vector Machine. This permits an automatic classification of the cell subpopulations selection in the bone marrow as well as the automatic counting of cells in each sample. The same process is applied for RBL, obtaining labeled leukemic B lymphocytes (LBL) depending on their maturation state. Labeling all cells means an improvement in the analysis of samples, since we can automatically distinguish cell subgroups and also perform more exhaustive studies on them.

As a second line of results, we consider the evolution of the disease over the available time points in each patient (Figure 8). It should be noted that the samples’ composition at diagnosis matches with the definition of B-ALL, as the B lymphocytes comprise around 80% of the whole bone marrow at that moment. Furthermore, the higher percentage of immature B cells in LBL is consistent with the idea of acute leukemia. This result leads to improved diagnosis in a single image, performing an automatic counting from raw flow cytometry data.

Beyond comparing ALL samples with RBM samples, the progression of the disease in the same patient has been observed using three time-point data: *t*_0_ (diagnosis), *t*_1_ (day +33 after the beginning of treatment) and *t*_2_ (day +78). This may provide the basis for new studies of disease progression as well as new models that predict whether a patient has a tendency not to respond to the treatment. In this sense, a particular topic to study is the difference between patients who relapse and patients who do not relapse, defined as R and NR patients respectively. We used statistical tools to identify whether there is a significant difference between those groups of patients over the three time points. The study of these data is approached from two perspectives, which involve calculating subpopulation percentages in each sample as well as determining differences for mean fluorescence intensity (MFI) in each sample’s immunophenotype. The same study was performed for B lymphocytes and their maturation states.

Our aim is the search for significant differences between the R and NR patients and compare them with RBM. We showed that there are no significant differences between the R and NR groups according to the percentage of the bone marrow subpopulations or their IPT markers MFI at diagnosis or day 33 of treatment. Nevertheless, all patients present differences with RBM cells at diagnosis. This fact aligns with leukemia diagnosis behavior, as it is the moment when the bone marrow is invaded by malignant B lymphocytes.

However, we have emphasized the significance of time *t*_2_, this is, day +78. This moment is of specific relevance in the patient’s follow-up, as it is the last routinely extracted bone marrow aspirate performed according to the SEHOP-PETHEMA 2013 protocol (see Section 2.3), when the treatment consolidation phase begins. Several results show that differences between R and NR patients which have been observed concerning this specific time point. For the study related to bone marrow composition and considering the percentages of B lymphocytes, R and NR differ in the percentage of B lymphocytes and erythroblasts. In fact, R patients’ B lymphocytes exhibit a behavior more similar to RBM than NR patients (Figure 9). This indicates that the two groups can be distinguished on day +78 of treatment focusing on study the amount of B lymphocytes. Also for the day +78, now in accordance to the MFI values of the IPT markers, the MWU test indicates that B lymphocytes CD19 of R patients show differences in comparison to NR patients. In Figure 10, it can be also observed that both for the CD19 and CD20 IPT markers, patients who relapse have MFI values more similar to RBM than patients who do not relapse. These markers are relevant as CD19 is an specific marker of B lymphocytes, and CD20 indicates differences in their maturation states. This result motivates a more thorough study now only for the LBL cells, where analogous percentage and MFI tests were performed. The results presented reinforced the idea that CD19 MFI is of significance when distinguishing R and NR at day +78, both for immature and more mature B lymphocyte maturation states. We developed a simple threshold-based classifier using CD19 and CD20 MFI values to distinguish patient groups, achieving moderate accuracy (up to 0.75, yet variable). To improve performance, we implemented a Random Forest model with oversampling and cross-validation, which slightly stabilized accuracy to 0.76. These results are shown in Tables 3 and 4. However, the limited sample size restricts generalization, highlighting the need for more diverse data in future studies.

Although our analysis reveals that the proposed pipeline shows considerable potential for assessing relapse risk at diagnosis in pediatric B-ALL, several limitations must be acknowledged. The retrospective nature of the study introduces potential biases and limits the standardization of sample collection and processing. Sample heterogeneity, both in clinical characteristics and disease stages, may affect subgroup consistency. Additionally, the restricted availability of cytometry markers constrains the depth of immunophenotypic profiling, which we have intended to overcome by means of sample merging. A clear limitation is the human subjectivity, which remains a factor in manual cytometry gating, despite efforts toward computational standardization. We have considered this by standardizing the data preprocessing, but again, batch effects introduce technical variability, such as differences in cytometer settings, which may also influence results. Regarding patient follow-up data, it can be lost due to changes in hospital care, affecting longitudinal analysis. Besides, clinical information, often extracted from medical reports, can be less quantifiable than cytometry data, making it more difficult to form homogeneous subgroups for reliable classification. Our samples, especially at early treatment time points like *t*_1_, exhibit low cellularity, which limits analytical resolution. While stratified subsampling to 10,000 cells per sample was implemented to ensure consistency, it inevitably results in the loss of cellular information, an issue that is particularly pronounced in low-cellularity samples. This, together with the dataset used, which comprises 234 samples from 75 patients, raises concerns about overfitting, particularly given the imbalance between relapsing (R) and non-relapsing (NR) patients. In fact, we were only able to train the regenerated patterns using the 9 samples from the day +78, as bone marrow extraction is a highly invasive procedure for pediatric patients. The available samples are those obtained due to suspected disease, which ultimately are considered to be healthy, hence the difficulty in acquiring more samples of this type. These limitations underscore the importance of ongoing efforts to validate the pipeline in a prospective study, which is currently underway and aims to address several of the issues outlined above.

The quantification of MRD by flow cytometry is based on the identification of cells that express the markers that define the leukemia-associated immunophenotype (LAIP), in the context of a hematopoiesis that is undergoing chemotherapy treatment. Our work deviates from the classic approach, that is, focused on the analysis of leukemic cells exclusively, to quantitatively study the regeneration of healthy hematopoiesis in the presence of insignificant MRD. The main contribution of this work is a novel, semi-automated computational pipeline (UMAP, DBSCAN, and SVM) that enables identification of leukemic cell subtypes using regeneration patterns from bone marrow samples, and tracks the kinetics of B-cell compartment recovery. This approach shows immunophenotypic patterns that distinguish patient cohorts and offer prognostic insight into relapse risk beyond standard MRD assessment. Our results demonstrate that this information has prognostic value, and suggest that the kinetics of regeneration of the B lymphoid compartment in the bone marrow are delayed in patients who subsequently relapse. The investigation cohort that these patients represent leads to results to be validated using an independent cohort, with the final aim of improving and implementing such computational tools in routine clinical settings.

**Supplementary information.** Supplemental information can be found online at https://github.com/ananinolopez/SI NinoLopez25.

## Declarations

### Ethics approval and consent to participate

This retrospective study was designed in accordance with the Declaration of Helsinki, under the LLAMAT Project protocol (2018). Approval was granted by the institutional review boards (IRB) of the Hospital Infantil Universitario Niño Jesús de Madrid. Patients and/or their legal guardians signed a written, formal informed consent to participate in the study. Personal information was anonymized.

### Consent for publication

Not applicable.

### Data, materials and code availability

The source code and functions used in this article can be consulted at https://github.com/ananinolopez/SI NinoLopez25. This repository also includes the full tables from the study of the patients and other figures related to iterations in the code.

### Author contribution

**ANL:** Conceptualization, Data curation, Formal analysis, Investigation, Methodology, Software, Writing—original draft, Writing—review & editing. **AMR**: Data curation, Investigation, Writing—review & editing. **RPG**: Data curation, Investigation, Methodology, Writing—review & editing. **ACR**: Data curation,Resources, Writing—review & editing. **MRO**: Conceptualization, Data curation, Resources, Writing—review & editing. **SC**: Conceptualization, Data curation, Investigation, Methodology, Writing—review & editing. **MR**: Conceptualization, Funding acquisition, Project administration, Supervision, Writing—review & editing

## Acknowledgements

This work was supported by project PID2022-140451OA-100 funded by Ministerio de Ciencia e Innovación/Agencia Estatal de investigación (doi:10.13039/501100011033). The support of Fundación Espanõla para la Ciencia y la Tecnología (FECYT project PR214), Asociación Pablo Ugarte (APU, Spain) and Junta de Andalucía (Spain) group FQM-201 is also acknowledged.

Editorial Policies for:

Springer journals and proceedings: https://www.springer.com/gp/editorial-policies

Nature Portfolio journals:

https://www.nature.com/nature-research/editorial-policies

*Scientific Reports*: https://www.nature.com/srep/journal-policies/editorial-policies BMC journals: https://www.biomedcentral.com/getpublished/editorial-policies

## Appendix A

### Abbreviations

Below is a list of abbreviations and acronyms used throughout this paper.

ALL: Acute Lymphoblastic Leukemia
BCa: Bias-Corrected and accelerate
CD: Cluster Differentiation
DBSCAN: Density-Based Spatial Clustering of Applications with Noise
FC: Flow Cytometry
HNJ: Hospital Niño Jesús
IPT: Immunophenotypic (marker)
IRB: Institutional Review Boards
LAIP: Leukemia-associated immunopheno-type
LBM: Leukemic Bone Marrow
LBL: Leukemic B lymphocyte
LLAMAT: Protocol: Early identification of resistance and optimization of treatments in childhood acute lymphoblastic leukemia through mathematical and statistical modeling
MFI: Mean fluorescence intensity
MRD: Minimal Residual Disease
MWU: Mann-Whitney U (test)
NR: Non-Relapse (patients)
R: Relapse (patients)
RBL: Regenerated B lymphocyte
RBM: Regenerated Bone Marrow
SEHOP-PETHEMA: ALL treatment protocol
SI: Supplementary Information
SVM: Support Vector Machine
UMAP: Uniform Manifold Approximation and Projection

